# Electrophysiological responses reveal a dedicated learning mechanism to process salient consonant sounds in human newborns

**DOI:** 10.1101/2024.09.06.611655

**Authors:** Paolo Barbieri, Pietro Sarasso, Alice Rossi-Sebastiano, Jacopo Frascaroli, Karol Poles, Chiara Peila, Alessandra Coscia, Francesca Garbarini, Irene Ronga

**Author notes:** **Corresponding author:** Irene Ronga, Department of Psychology, University of Turin Via Verdi 10, 10124 Turin, Italy. **Author contributions: PB:** Designed research, Performed research, Analyzed data, Wrote the paper; **PS:** Designed research, Performed research, Analyzed data, Wrote the paper; **ARS:** Performed research, Analyzed data, Wrote the paper; **JF:** Designed research, Performed research, Wrote the paper; **KP:** Performed research, Analyzed data, Wrote the paper; **CP:** Designed research, Performed research, Wrote the paper; **AC:** Designed research, Performed research, Wrote the paper; **FG:** Designed research, Analyzed data, Wrote the paper; **IR:** Designed research, Performed research, Analyzed data, Wrote the paper. **Competing interest statement:** The authors declare no competing interests.

## Abstract

Isolating relevant sounds in the auditory stream is a crucial feature accomplished by human infants and a pivotal ability for language acquisition. Therefore, it is reasonable to postulate the existence of early mechanisms reorienting attention toward salient acoustic stimuli. Previous studies suggest that infants consider consonant sounds as more salient than dissonant ones, because the former resemble human vocalizations. However, systematic evidence investigating the neural processes underlying consonance tuning in newborns is still scarce. Here, we investigate newborns’ ability to recognize and learn salient auditory stimuli by collecting Mismatch Responses (MMRs) to consonant and dissonant sounds and by computing the trial-by-trial correlation of the neural signal with Bayesian Surprise (a theoretical measure of learning). We present 22 healthy newborns (40.4 ± 15.8 hours) with a pseudo-random sequence of deviant and standard auditory events, while we record their electroencephalogram. Our results show that newborns exhibit a neural encoding of auditory regularities for all sound types (consonant and dissonant), as demonstrated by the presence of MMRs and significant correlation of the neural signal with Bayesian Surprise. Furthermore, consonant and dissonant sounds elicited MMRs and correlations with Bayesian Surprise of opposite polarities, with consonant auditory stimulation evoking negative responses, reminiscent of an adult-like MMR. Overall, our findings suggest that newborns display a dedicated perceptual learning mechanism for salient consonant sounds. We speculate that this mechanism might represent an evolutionary-achieved neural tuning to detect and learn salient auditory stimuli with acoustic features resembling human vocalizations.

**SIGNIFICANCE STATEMENT:** Discriminating salient sounds in noisy sensory streams is a fundamental ability displayed by human infants, pivotal for acquiring crucial skills including language. Our study shed light on this ability by: (1) investigating perceptual learning mechanisms in newborns’ with a neurocomputational approach; (2) exploring the role of salient consonant sounds in modulating such mechanisms. Since human vocalizations are often consonant, the presence of a mechanism dedicated to enhance the processing of consonant sounds in newborns would confer evolutionary advantages. Our findings, indicating that newborns possess a dedicated and more refined perceptual learning mechanism to process consonance, corroborates this hypothesis. We speculate that this neural mechanism might facilitate the identification of salient acoustic input and support language acquisition in early infancy.

## Introduction

The ability to discriminate and track salient sounds while being immersed in a rich and noisy sensory environment underpins many human abilities, including linguistic communication^1,2^. One should therefore expect to find neural traces of the development of such a crucial ability early on in the lifespan.

Although findings are still controversial^3^, preliminary behavioural evidence suggests that newborns and infants seem to consider consonant sounds as more salient than dissonant ones, since they tend to allocate more attention to the former^4–10^. This finding is often interpreted in light of the *Vocal Similarity Theory*, which assumes that humans are provided with a specific mechanism directed to enhance the processing of sounds similar to the human voice (which is mainly consonant), as this would favour the acquisition of communication skills^4,11^. Consistently with this hypothesis, in a recent study on adults, we found that enhanced implicit learning mechanisms are observed for preferred, consonant sounds, as demonstrated by the magnification of the Mismatch Negativity^12^. The Mismatch Negativity (a common mismatch response obtained by subtracting the responses evoked by standard, repeated stimuli from those evoked by deviant ones) is considered a well-validated neurobiological marker of implicit learning, as well as an important window into how we keep track of sensory regularities and learn from relevant, most informative stimuli^13,14^. Unlike adults, newborns present two distinct mismatch responses (from now on, MMRs) with opposite polarities^15^. Deviancy detection can elicit positive^15^, and (more rarely) negative MMRs^1,16–18^, sharing similar latencies (between 0.25 and 0.45 s) and scalp distributions (fronto-central). Interestingly, negative MMRs are usually associated with later stages of development^15,19–21^ and with the detection of native language stimuli, even in the first year of life^16,18,22^. However, whether the early polarity switch of the MMRs may be related to the processing of more relevant consonant sounds is still unclear. Here we aim to test whether MMRs’ polarity switch observed in newborns actually represents an early tool for differentiating consonant and dissonant stimuli, with the final aim of detecting the most informative stimuli in the entropic sensory context.

In the present electrophysiological study, we investigate newborns’ ability to track and discriminate informative sounds in the sensory stream by examining their MMRs with a roving paradigm^23,24^. In a roving paradigm, participants are presented with a sequence of standard, repeated tones interrupted by pitch-deviant ones^12,25^. Unlike in classical oddball paradigms, in roving paradigms each auditory stimulus (either high- or low-pitch intervals in our experiments) has the same probability of occurrence and can represent both a deviant and a standard event, depending on its position in the stimulus sequence^24–27^ (see Figure 1). In this way, the roving paradigm allows to differentiate deviancy responses from a generic rarity effect that could occur in sequences with an unmatched probability of occurrence between different stimuli.

**Figure 1.**
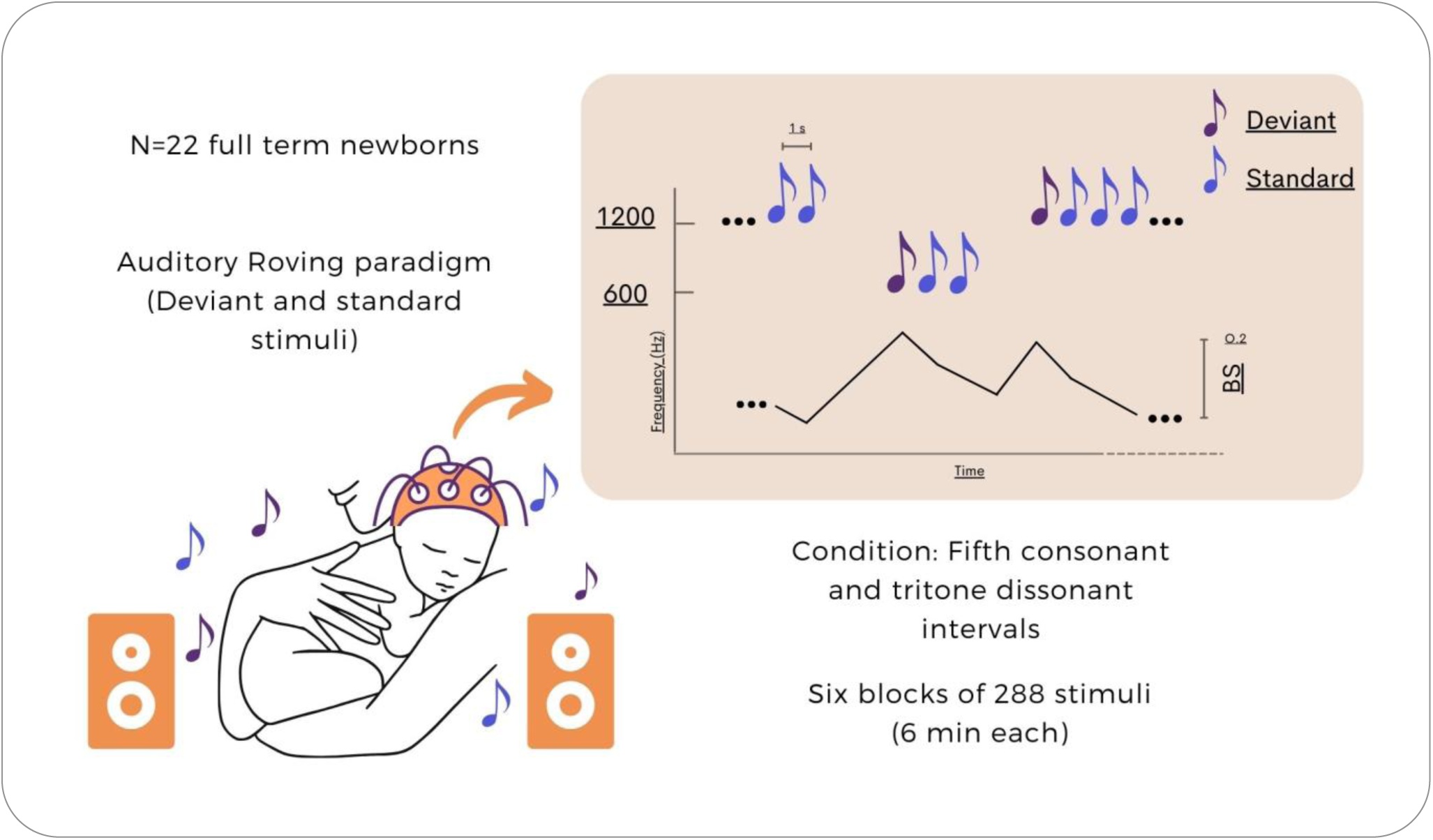
In our roving paradigm, newborns listened to six blocks of 288 sounds. We presented fifth (consonant) and tritone (dissonant) sound intervals separately: three blocks were composed of sequences of high pitch and low-pitch consonant intervals, while the other three were composed of sequences of high pitch and low-pitch dissonant intervals. The top-right graph represents an example of a sequence of standard and deviant sounds. The frequency (Hz) is represented on the y-axis. Sound intervals that differed from the preceding one were considered deviant.

The adoption of a roving paradigm in the present experiment allows us to address two important questions about early auditory porcessing. First, we explore how newborns encode sensory regularities and update these encodings in light of surprising sensory inputs (a process that from now on we will call *perceptual learning: see*^28^). Second, we assess whether a specific perceptual learning mechanism, directed to process consonant sounds, is already present in the first hours of life. With these aims in mind, we also perform a trial-by-trial correlation between the brain signal elicited by each auditory event in the roving sequence and the values obtained with a Bayesian Surprise algorithm. Bayesian Surprise is a mathematical measure of how much new information is conveyed by each stimulus in a sequence^25,29^. The correlation of the neural signal with this mathematical measure describes the degree to which the representations of the sensory environment are updated in response to deviant, information-rich stimuli^30^. In other words, this correlation can be considered an index of how effectively we are learning from the environment. Interestingly, in adults, the amplitude of Mismatch Negativity correlates with the trial-by-trial fluctuations of Bayesian Surprise, thus confirming that MMRs reflect perceptual learning dynamics in adults^12,31,32,32^. However, it is still unknown whether this correspondence between MMRs and trial-by-trial fluctuations of Bayesian Surprise is already present at birth and, in case it is, whether it is similar for the processing of consonant and dissonant sounds.

In sum, in the present study we aim to: (1) investigate newborns’ perceptual learning of auditory regularities in a roving paradigm, and test whether the brain signal elicited by each auditory event in the sequence correlates with Bayesian Surprise values; (2) verify the presence of a dedicated perceptual learning mechanism to process novel consonant *vs.* dissonant stimuli, as measured by differences in MMRs and correlation with Bayesian Surprise. As results, (1) we expect to observe MMRs to both consonant and dissonant intervals and a significant correlation between the trial-by-trial fluctuation of the neural signal and Bayesian Surprise values. (2) We predict that, in line with the Vocal Similarity Theory, MMRs in response to consonant sounds will be significantly different from the MMRs elicited by dissonant sounds, providing newborns with a a neural mechanism able to discriminate salient consonat stimuli, similar to the human voice spectrum.

## Results

### Perceptual learning in newborns

*Response to standard and deviant events.* Newborns’ (mean age ± SD: 40.4 ± 15.8 hours since birth) neural responses elicited by consonant (fifth) and dissonant (tritone) intervals recorded from Fpz are reported in Figure 2A. The grand-average (N = 22) waveforms were consistent with those observed in previous studies on newborns, highlighting distinct responses to deviant *vs.* standard stimuli, within the time window 0.2 - 0.45 s, over fronto-central electrodes^1,15,17,21,22^ (Figure 2A). Interestingly, larger negative waveforms emerged in response to consonant deviant as compared to consonant standard stimuli, whereas dissonant deviant stimuli elicited greater positive waveforms as compared to dissonant standard ones (Figure 2, Panel A). Accordingly, MMRs were elicited by both consonant and dissonant intervals, and showed two peaks of opposite polarity at approximately 0.2 and 0.4 s post-onset (Figure 2C, MMR), thus indicating the ability of newborns to recognize environmental auditory regularities (i.e., discriminating deviant events from standard events) in a pseudo-random stream of sensory events^1,15,17,21,22^.

**Figure 2.**
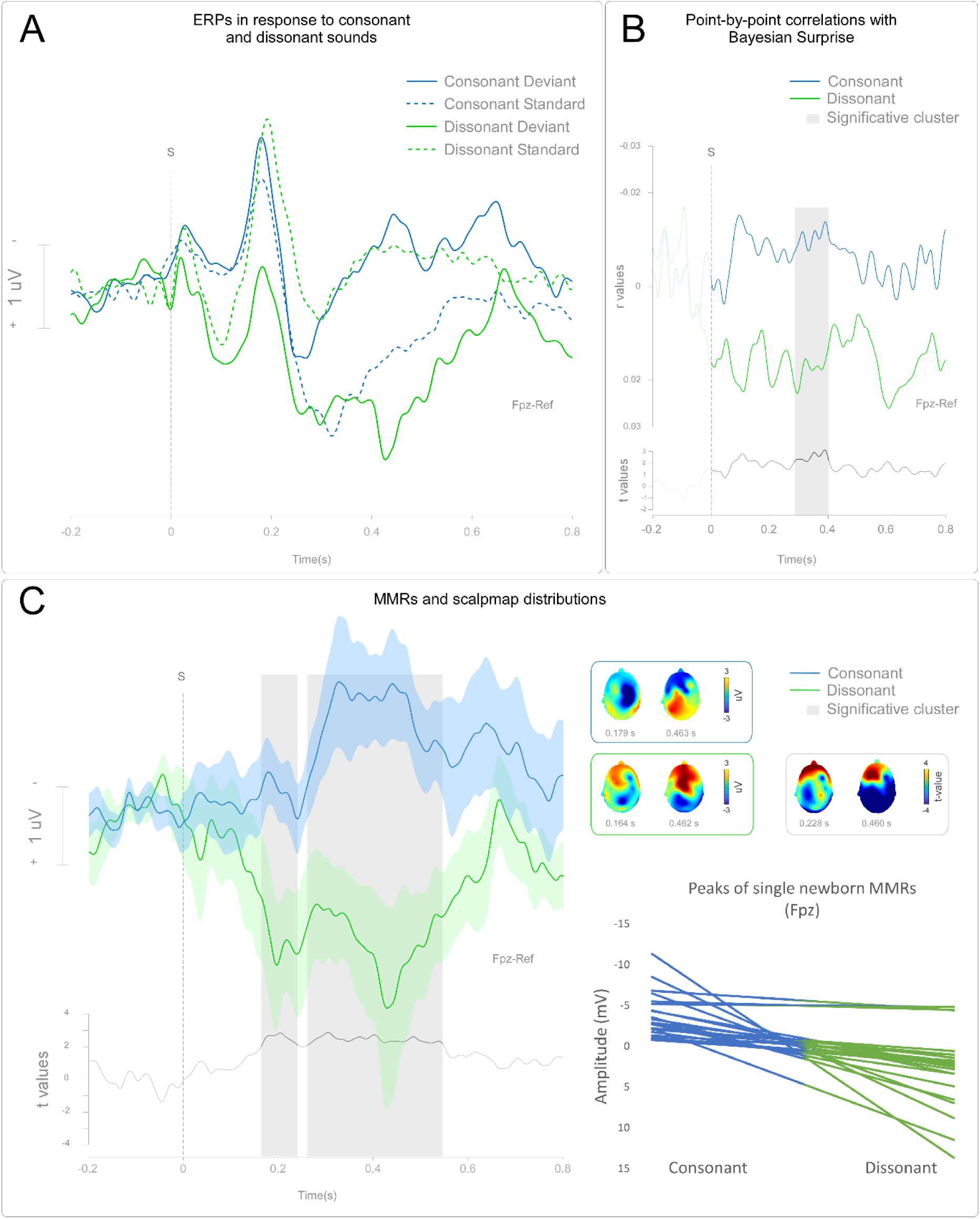
*Panel A.* Grand-average ERP registered from Fpz for consonant and dissonant sounds. X axis: time (s); Y axis: amplitude (uV). S = stimulus onset. *Panel B.* The waveforms represent grand-average r values between trial-by-trial neural signal fluctuations registered at Fpz and Bayesian surprise. Shaded areas highlight significant time clusters between conditions (point-by-point statistics). *Panel C.* Grand-average MMR waveforms for consonant and and dissonant sounds registered on Fpz. Shaded areas highlight significant time clusters between conditions (point-by-point statistics). The scalpmaps on the top-right panels show the scalp distribution of MMRs around 0.2 and 0.4 s post-onset and the distribution of the significant difference between MMRs to consonant and dissonant sounds around 0.2 and 0.4 s post-onset. The bottom-right graph shows the peaks (time window: 0.25-0.4 s) of each newborn’s MMRs to consonant and dissonant sounds.

#### Point-by-point correlations with Bayesian Surprise

Four newborns were excluded from the sample of the Bayesian surprise analysis, since we were not able to collect enough trials during the experiment to perform the trial-by-trial analysis (due to deep sleep or crying). The outcome of the point-by-point correlation between trial-by-trial neural signal amplitudes and Bayesian values (final sample N=18) returned 1 s-long (from 0.2 s pre-onset to 0.8 s post-onset) time series of r values for each channel and condition (consonant *vs.* dissonant intervals). The results indicate that r values (with specular polarity for consonant and dissonant intervals; see Figure 2, panel B) peaked at around 0.1 s, 0.3 s, and 0.4 s post-onset, which corresponds to the MMR time window. When exploring the mean amplitude (0.2-0.45 s post-onset) of the r values time series elicited by consonant and dissonant intervals, we found that they were both significantly different from the constant zero (consonant vs 0, T = −2.00, M = −0.006, SD = 0.03, p = 0.04; dissonant vs 0, T=3.42, M = 0.01, SD = 0.008, p = 0.003). Similar results were obtained when exploring the r values time series with a point-by-point t-test. The time series elicited by consonant intervals were significantly different from zero in the time window between 0.1 and 0.4 s post-onset over frontoparietal channel (F4 and P3), while the r values time series elicited by dissonant intervals were significantly different in the time window around 0.6 s over frontocentral channel (Fpz, F7, F3, Fc1 and Cz). Overall, these results are in line with our previous studies on adults^12^, showing that MMR amplitude fluctuations significantly correlate with Bayesian Surprise values. This finding seems to indicate that, in newborns too, MMR amplitude indexes perceptual learning.

### MMRs to consonant and dissonant sounds

*MMRs*. To further investigate the MMRs elicited by consonant and dissonant intervals, we performed whole-brain point-by-point t-tests, and, in accordance with previous newborn studies (see § Methods), we compared the ERP mean amplitude between 0.2-0.45 s over frontal electrodes (see § Methods). Importantly, both analyses yielded comparable results. Consonant intervals elicited a negative MMR, while dissonant intervals elicited a positive MMR (Figure 2C). More specifically, the subtraction of standard intervals and deviant ones (i.e., the MMR) reflects the opposite polarity of the ERP waveforms described above, generating two MMR with specular polarity in response to consonance and dissonance.

Both the two-tail paired sample t-test performed on mean amplitude (T = −2.22, M = −1.49, SD = 0.67, p = 0.028) and the whole brain point-by-point t-test revealed that the MMR waveforms elicited by consonant and dissonant intervals were significantly different (Figure 2, Panel C). The point-by-point statistics revealed two significant time windows, between 0.165 and 0.245 s (corresponding to MMR first peak), and between 0.26 and 0.546 s post-onset (corresponding to MMR second peak). Interestingly, also the scalp topographies of the MMRs elicited by consonant and dissonant stimuli appeared different (Figure 2, panel C). More specifically, the MMRs elicited by consonant stimuli display a broad frontocentral negativity, with a lateralized (right hemisphere) activity peaking at 0.2 s, and a left-hemisphere lateralization peaking at 0.4 s. On the other hand, MMRs elicited by dissonant stimuli display a broad and less lateralized positive, fronto-central activity, peaking at 0.2 and 0.4 s.

#### Point-by-point correlations with Bayesian Surprise

Both the paired sample t-test on mean amplitude (i.e., T = −4.09, M = - 0.017; SD = 0.004; P < 0.001) and the point-by-point t-test revealed that the r values waveform elicited by consonant and dissonant intervals were significantly different in the time interval between 0.1 and 0.4 s (corresponding to the latency of MMRs) over frontocentral electrodes (Fpz, Fp2, F4, Fc1, Fc2 – see Figure 2B).

## Discussion

In the present experiment, we explored newborns’ perceptual learning dynamics with an auditory roving paradigm. Our results show that, as early as a few hours after birth, newborns exhibit a neural encoding of the regularities in their auditory stream for all the presented sound types (consonant and dissonant), as demonstrated by the presence of MMRs and by the trial-by-trial correlation with Bayesian surprise values. Furthermore, our EEG results showed that newborns display two different patterns of neural response for consonant and dissonant sounds, supporting the hypothesis that two distinct neural mechanisms may be involved in their processing. We speculate that such neural specialisation might represent an evolutionary-achieved neural tuning to detect and discriminate salient auditory stimuli, with acoustic features resembling human voice spectra.

### Auditory perceptual learning in newborns

Overall, our findings highlight newborns’ ability to generate and update encodings of auditory regularities according to the informativeness of each new event in the unfolding auditory stream. Previous research exploiting oddball paradigms or other repetition-suppression tasks (where responses to frequent stimulation are compared to the responses evoked by rare events) showed that newborns are able to discriminate stimuli in different sensory modalities within hours of birth^17,33,34^ and even during intrauterine life^35–37^. For example, Kushnerenko and colleagues showed that newborns exposed to an oddball task were able to discriminate a variety of auditory features, such as pitch and duration^38^. Here, we demonstrate that even when the probability of occurrence of each stimulus in the sequence is matched (as in the present roving paradigm), newborns still display MMRs, which show their ability to detect random, local violations of a pattern. Furthermore, we showed that the amplitude of trial-by-trial responses to auditory events significantly correlates with Bayesian Surprise values, similarly to what is observed in adults^12^. This finding indicates that the amplitude of MMRs in newborns is directly related to a mathematical measure (i.e., Bayesian Surprise) of the amount of novel information carried by each stimulus in the auditory sequence.

In a way, it is not surprising that an effective encoding of regularities and deviations in the auditory stream can be observed a few hours after birth, given the role that this encoding plays in the earliest human developmental stages. Among other things, this encoding might help to readily build expectations about different auditory features and to discriminate the most informative stimuli (such as distinguishing language sounds from other auditory stimuli^15,33,39^).

### An early mechanism to detect change in salient consonant sounds?

Consonant and dissonant intervals elicited MMRs and point-by-point correlations with Bayesian Surprise of opposite polarities, with consonant auditory stimulation triggering negative MMRs, reminiscent of an adult-like mismatch negativity^12,13,31^. Importantly, even though deviancy detection can be observed for both consonant and dissonant sounds (see Figure 2), deviant consonant stimuli generate more negative responses than standard ones, whereas deviant dissonant sounds elicit more positive responses than standard ones (Figure 2, panel A). This finding suggests the presence of dissociable neural mechanisms dedicated to the processing of consonant *vs*. dissonant stimuli. This idea is further supported by the scalp distribution of the MMRs. Negative MMRs show a negative polarity over frontal electrodes and a positive polarity over central electrodes. Conversely, positive MMRs present an opposite distribution (see Figure 2, panel C). Negative MMRs are usually associated with later developmental stages, and a transition from positive to adult-like, negative MMRs can be gradually observed within the first year of life^15,19,40^. Interestingly, our results seem to indicate that, already a few hours after birth, a distinct and more adult-like perceptual learning process can be observed for consonant sounds only. This result is in line with the findings of a previous study using an oddball paradigm^17^, which showed that newborns’ electrophysiological responses are different for different types of deviant chords (such as a minor or dissonant chord included in a stream of major chords). In the present study, by employing a roving paradigm (which, as we said, allows to differentiate between novelty and rarity effects), and through a neurocomputational approach based on Bayesian statistics, we provide compelling evidence that newborns present distinct mechanisms for the processing of consonant and dissonant intervals.

Previous behavioural research on gaze orientation^9^ and sound production^10^ seems to support the hypothesis of a preferential processing for consonant sounds in newborns. More specifically, Masataka shows that newborns looked longer towards objects producing consonant compared to dissonant sounds. Furthermore, Di Stefano and colleagues (2017) showed that newborns prefer to generate consonant rather than dissonant harmonic intervals when interacting with a musical tool. Our electrophysiological findings may shed light on the neural underpinnings of these behavioural effects. It seems plausible that the adult-like perceptual learning process observed for consonant stimuli may support the distinction between consonant and dissonant sounds in the early infancy, and be evidence that more attentional resources are allocated to former rather than the latter right from birth. Such an early dissociation in the processing of consonant and dissonant sounds might indicate the presence of an evolutionary advantage granted to newborns that are able to better discriminate consonant auditory stimuli.

But what might be the adaptive value of a neural tuning for consonance at birth? As we have anticipated, according to the Vocal Similarity Theory, the preference for consonance is explained by the similarity to the human voice spectrum^4^. It is possible that, throughout evolution, consonant stimuli resembling human vocalizations acquired a distinct perceptual learning process advantageous for language acquisition^4^. The idea of a connection between language development and advanced perceptual learning phenomena seems to be confirmed by other electrophysiological studies on young infants. The effectiveness of auditory perceptual learning in the first months of life has been shown to be a crucial prerequisite for later language development^1,22,41^. Friedrich and colleagues (2009) demonstrated that children displaying age-appropriate expressive language abilities at 2.5 years exhibited an early (at 3-4 months) negative MMR when processing native language deviant stimuli in an oddball design. Conversely, infants developing poorer language skills did not display early negative MMRs^1^. Similarly, Garcia-Sierra and colleagues (2016) demonstrated that the quantity of linguistic input infants receive during their first year of life predicts neural perceptual learning maturity, as indexed by the presence of negative MMRs^22^.

The difference in the processing of consonant stimuli that we observed, therefore, might be the product of a more refined neural mechanism possibly developed for early language recognition^37^. Dissonant stimuli, on the other hand, could be elaborated by a more basic pathway, devoted to general, less salient, acoustic input^41^. An extensive literature confirms that humans, even during intrauterine life, discriminate voice and speech sound and display an early left-hemispheric specialization for linguistic processing^36,37^. Interestingly, in the present study too, the scalp distribution of the MMRs elicited by consonant intervals displays a left lateralization over frontocentral electrodes, whereas dissonant intervals elicited a more widespread topographical distribution in the parieto-central electrodes (Figure 2; Panel C). If further confirmed by future studies exploring the source localization of MMRs, this finding might therefore be considered as supporting evidence for the involvement of a primitive language-related pathway in consonant stimulus processing.

### Nature or Nurture?

Our study could contribute to the ongoing discussion about the respective roles of biology (nature) and cultural exposure (nurture) in determining consonance and dissonance perception^4^, an issue strongly debated in musicology and auditory psychology and neuroscience.

According to the ‘nature’ perspective, consonance preference has generally been interpreted within two different frameworks: the Vocal Similarity Theory^4^, which we discussed above, and the Peripheral Origin of Consonance Discrimination^42,43^. While the former emphasizes the similarity between consonant stimuli and the human vocal spectrum, the latter highlights the role of mathematical proportions of chords and their interaction with the auditory system (i.e., basilar membrane, auditory nerve and brainstem), in explaining consonance preferential processing^42,43^. Our results, indicating an early tuning for consonance sensory processing, seem to lend support to the ‘nature’ explanation. Indeed, it seems evolutionary adaptive to display a preferential neural pathway to process stimuli belonging to one’s own species’ vocalizations.

On the other hand, the ‘cultural’ perspective usually emphasized competence acquired through exposure to the Western music system^44^. Our previous study on adults showed that 1 out of 5 participants displayed an identical neural tuning (and aesthetic preference) to dissonant sounds, with a magnification of the Mismatch Negativity for these stimuli^12,31,32^. We speculated that such a neural tuning for dissonant stimuli, rather than innate, was gradually developed in response to specific life experiences^44–50^. Crucially, the ‘nature’ and ‘nurture’ hypotheses are not necessarily mutually exclusive: in all likelihood, biological and cultural perspectives should be used synergically to explain the complexity of consonance and dissonance perception.

To sum up, we speculate that at birth newborns are provided with a preferential neural pathway to process consonant input. This might be due to the similarity between human vocal stimuli and consonant sounds, which makes them more salient as compared to dissonant sounds. At a later developmental stage, experience could shape and refine this process and perhaps tune infants, for example, toward native language or phonologically relevant information. Throughout development, adults might eventually devote this preferential pathway (signaled by magnified mismatch negativity responses) to stimuli which have come to be considered more salient, either by experience or by nature. It is worth noting that neural tuning in adults is systematically associated with an aesthetic reward^31,51^, since greater MMRs are consistently correlated with subjective aesthetic preferences^12,31,32^. Future studies might therefore explore the emergence and development of an explicit aesthetic preference throughout infancy.

### Conclusions and future directions

In conclusion, our results suggest that newborns exhibit electrophysiological correlates of auditory perceptual learning processes within hours of birth and that these processes differ for consonant and dissonant stimuli. A specific neural process seems to be dedicated to the processing of novel, consonant stimuli, providing newborns with an early mechanism to discriminate salient information in the noisy auditory environment, likely facilitating language acquisition.

Further evidence will be needed to confirm this hypothesis and further clarify perceptual learning processes throughout infancy. More specifically, future studies should be directed to further investigate the neural encoding of sensory regularities in newborns and other possible factors that might modulate it. As an example, it may be important to investigate the role of different physical and contextual features, such as the presence vs. absence of social interaction, in modulating perceptual learning. Finally, it would be interesting to investigate how such neural mechanisms emerge within ontogeny from the intrauterine life to early infancy.

## Methods

### Participants

In the present experiment, we recorded the electroencephalography (EEG) of 22 newborns (mean age of 40.4 ± 15.8 hours at the time of testing) to compute event-related potentials (ERPs) in response to an auditory roving paradigm^25^. Parents provided written informed consent and the Ethical Committee of Sant’Anna University Hospital approved the study (n. 0121061; 14/12/2017-14/12/2022). The original sample size (N = 25) was a priori determined so as to match the average number of participants involved in previous ERP studies on newborns^18,52^ and in a very similar paradigm on adults^12^. Three newborns were discarded due to poor signal-to-noise ratio (mostly because of body movements during the EEG recording). Newborns included in the final dataset (n=22) were awake during the presentation of the experimental conditions, and relaxed in their parent’s (mother or father) arms.

### Data availability

Raw data used in the study and the tables containing the mean amplitude used in the analysis will be made publicly available upon acceptance for publication.

### Stimuli and Procedure

Newborns passively listened to a standard roving paradigm (overall recording duration: from 30 minutes to 1 hour, depending on the newborn’s compliance), while we registered their neural activity with 30-channel electroencephalography (EEG). Newborns listened to three sequences of consonant sounds (fifth intervals) and three sequences of dissonant sounds (tritone intervals), presented in blocks of 6 minutes each (with trains of 288 stimuli per block). Sounds (50 ms duration) of alternating high/low pitch were presented at a frequency of 1Hz (inter-trial interval: 1 s). Note that all sounds included in the sequence had the same probability of occurrence (Figure 1). For further detail about the stimuli and the apparatus please refer to Supplementary Information.

### EEG preprocessing and statistical analysis

#### EEG preprocessing

An extended version of EEG analyses is reported in the Supplementary Information. Continuous EEG data were segmented into epochs including a 0.2 s before stimulation and 0.8 s after stimulation (total epoch duration: 1s), and band-pass filtered (1-30 Hz) using a fast Fourier transform filter (in accordance with previous literature exploring ERP responses in newborns^52^ and MMN effects and Bayesian surprise in adults^12,25^. Each epoch was baseline corrected using the interval from −0.2 to 0 s. Artifacts due to eye blinks or eye movements were eliminated using a validated method based on an Independent Component Analysis (ICA ^53^) while noisy epochs were discarded via artifact rejection. Epochs belonging to the same interval type (i.e., consonant or dissonant) and to the same condition (i.e., standard vs. deviant) were averaged time-locked to the onset of the stimulus, thus yielding four average waveforms (Consonant Deviant, Consonant Standard, Dissonant Deviand, Dissonant Standard) for each subject. To compute MMRs, the responses to standard-repeated sounds were subtracted to the response to deviant sounds^13^ for each newborn and for each condition (consonant vs. dissonant) separately. Furthermore, we computed trial-by-trial Bayesian surprise values, indexing the informative value conveyed by each single trial/sound included in the sequence (for a total of 1728 Bayesian surprise values). Bayesian Surprise Index was computed as the Kullback-Leibler divergence between prior and posterior distribution using a sequential Bayesian learning algorithm of stimulus probabilities^12,24,25,54,55^.

#### Statistical analyses on ERPs

To compare our experimental conditions, we employed two different statistical approaches. First, we performed a whole-brain, fully data-driven analysis, without any a priori assumption, by computing point-by-point t-tests to compare single subjects’ waveforms, at each time point and for each channel separately^12,31,56,57^. To control for multiple comparisons, we then performed a clustersize-based permutation correction based on temporal consecutivity and spatial adjacency (1,000 permutations; alpha level = .05; percentile of mean cluster sum = 95; minimum number of adjacent channels = 3). Furthermore, we employed a second statistical approach widely used in newborns studies^52^, comparing the mean amplitude recorded in a time window and scalp distribution of interest. For the present research, we selected a time window between 0.2 and 0.45 s and a fronto-central region of interest (see Supplementary Information for further detail), since MMRs in newborns are usually recorded at these latency and scalp distribution^1,2,15–18,21,22^.

Therefore, we performed a point-by-point t-test and a paired-sample t-test (with mean amplitude values) to compare single subjects’ MMR amplitudes to consonant and dissonant sounds. Finally, we also computed a 2 x 2 Anova on the average ERP of each participant to further clarify the role of *consonance* and *deviancy* factors and their interaction. The analysis and results are extensively described in the Supplementary Informations and fully confirmed the analyses on MMRs.

#### Correlation with Bayesian Surprise values

To compute the correlation with Bayesian Surprise, we performed a point-by-point correlation analysis between single-trial amplitude fluctuations (i.e. 864 epochs for each condition) and Bayesian Surprise values corresponding to outputs of Bayesian Surprise algorithm^12,51,58^. R values constituted the input for the subsequent group-level analyses. As a first step, we performed a point-by-point t-test on each r-value time series (consonant and dissonant separately) against the constant zero. Furthermore, we performed one t-test against the constant zero for each condition separately, employing the mean amplitude of the r-value time-series between 0.2 and 0.45 s (region of interest: frontocentral electrodes). These analyses were crucial to verify that the r-value time series of consonance and dissonance conditions were significantly different from zero, thus indicating that the neural signal fluctuations actually reflected a form of Bayesian learning updating. Furthermore, to verify whether the correlation with Bayesian Surprise significantly differed across consonance conditions, we performed a two-tailed point-by-point t-test comparing single subjects correlation coefficients, for each condition (consonance vs. dissonance) as well as a standard t-test, comparing consonant and dissonant condition and employing the mean amplitude of the r-value time-series between 0.2 and 0.45 s (region of interest: frontocentral electrodes).

## Acknowledgements

The Authors are grateful to the participants involved in the present study. **IR** is funded by PON project “Ricerca e Innovazione” 2014-2020 (Italian Ministry of University and Research; D11B21005800007); **JF** is founded by the project NODES “Nord ovest digitale e sostenibile” of the PNRR (Unione Europea - Next Generation EU: D17G22000150001); **FG** is founded by the European Union (ERC-STG, MyFirstBody, 101078497). The views and opinions expressed are however those of the authors only and do not necessarily reflect those of the European Union or the European Research Council Executive Agency. Neither the European Union nor the granting authority can be held responsible for them.

## Supporting Information

### A Method

#### A1 Participants

22 full term healthy newborns (females: 10; gestational age > 37 weeks) participated in the Experiment conducted at the Sant’Anna University Hospital, Città della Salute e della Scienza di Torino, Italy, where they were born. Inclusion criteria were identical to those of the the American College of Obstetricians and Gynecologists (ACOG), i.e., birth weight > 2500 g, weeks, and Apgar index score = 9 at 5 minutes of life. No evident abnormalities were recorded at birth in the participants. The 22 newborns included in the final sample had a mean age of 40.4 hours at the time of testing (SD = 15.8), a mean birth weight of 3.267 g (SD = 361), and a mean gestational age at birth of 38,9 weeks and 3 days.

All newborns’ parents were fully informed about the experimental procedures prior to the experiment and they gave their written informed consent to allow the study. The study conformed to the standards required by the Declaration of Helsinki and was approved by the local ethics committee (University of Turin; prot. N° 0121061). The original sample size (N = 25) was a priori determined to match the average number of participants involved in previous ERP studies on newborns ^52^. Three newborns were discarded due to poor signal-to-noise ratio (mostly because of body movements during the EEG recording). Newborns included in the final dataset (n=22) were awake, relaxed in their mothers’ arms, who were comfortably sitting on an armchair. The experiment could last between 30 minutes and an hour for the situational needs of newborns, indeed we aimed at testing them when they were awake and relaxed, to wait for the best moment to collect the data.

#### A2 Stimuli and Apparatus

Intervals were created with Csound () which allows to select the frequency of the two notes composing the interval, which were played simultaneously for 50 ms via loudspeaker. Loudness of sounds was kept equal across subjects and set at a comfortable level (64 dB). The ratio between the frequency of the two composing notes defined two interval types (i.e. Fifth consonant and Tritone dissonant): the smaller the integer numbers that define the ratio, the more consonant will be the interval ^59,60^. Perfect fifth intervals, usually perceived as more consonant by adults, have a ratio of 3:2, while tritones, frequently categorised as more dissonant are defined by a ratio of 45:32. For each interval type (fifths or tritones), we employed two different pitches (i.e., high vs low). Low-pitch and high-pitch fifth intervals were composed of notes with a frequency of 150 and 100 Hz and 600 and 400 Hz, respectively. Low-pitch and high-pitch tritone intervals were composed of notes with a frequency of 150 and 106 Hz and 600 and 426 Hz, respectively. During the experiment, newborns were relaxed on their mothers’ lap and the loudspeaker distanced 20 cm from them.

#### A3 Electroencephalogram recording and processing

Newborns electrophysiological neural responses to the auditory roving paradigm were recorded using 32 Ag-AgCl electrodes placed on the scalp according to the International 10-20 system and referenced to the nose. Electrode impedances were kept below 5 kΩ. The electro-oculogram (EOG) was recorded from two surface electrodes, one placed over the right lower eyelid and the other placed lateral to the outer canthus of the right eye. Signals were digitized at a sampling rate of 2048 Hz with a Handy EEG amplifier (SystemPlus Evolution, Micromed, Treviso, Italy). EEG data were offline pre-processed and analysed using Matlab (Mathworks, Natick, MA). Continuous EEG data were segmented into epochs including a 0.2 s before stimulation and 0.8 s after stimulation (total epoch duration: 1s), and band-pass filtered (1-30 Hz) using a fast Fourier transform filter (in accordance with previous literature exploring ERP responses in newborns and MMN and Bayesian surprise in adults ^12,25,52^). Each epoch was baseline corrected using the interval from −0.2 to 0 s from the stimulus trigger. Artifacts due to eye blinks or eye movements were eliminated using a validated method based on an Independent Component Analysis (ICA ^61^) while noisy epochs were discarded via artifact rejection. Epochs belonging to the same interval type (i.e., fifth or tritone) and to the same condition (i.e., standard vs. deviant) were averaged time-locked to the onset of the stimulus trigger, thus yielding four average waveforms (Fifth Deviant, Fifth Standard, Tritone Deviand, Tritone Standard) for each subject.

#### A4 Roving paradigm and Procedure

Newborns passively listened to six runs of a standard roving paradigm with trains of 288 stimuli per run (overall recording duration: from 30 minutes to 1 hour, depending on the newborn’s compliance), while we registered their neural activity with electroencephalography (EEG). Three runs of only fifth consonant (high-pitch and low-pitch) and three runs of only tritone dissonant (high-pitch and low-pitch) intervals were presented (the number of runs could also differ depending on the newborns’ compliance). The order of presentation of fifth and tritone runs was randomised across participants, to exclude any specific sequence effects. In roving paradigms, in contrast to traditional oddball ^62^, different stimuli (high-pitch and low pitch in our experiment) represent both Deviant and Standard stimuli ^12,25–27^, and each stimulus has exactly the same probability to occur, thus allowing to dissect genuine effects of Bayesian perceptual learning from rarity-driven modulations ^12^. During the experiment, newborns were presented consecutive trains of high and low pitch intervals with a constant interstimulus interval of 1 s ^12^; any time a particular stimulus (low pitch or high pitch) repeat itself constitutes a sequence of Standard stimuli while when the stream of standard sound was interrupted by a different stimulus (low or high pitch), the first stimulus of the new train constitutes a Deviant event, since it differs (by its pitch) from the preceding train of Standard stimuli ^12,63^. Similarly, the repetition of the Deviant stimulus constitutes a new sequence of standard, repeated, intervals. A pseudorandom order was used to constitute the length of the train of high and low pitch interval, so that both the number of presentation and the average value of the Bayesian surprise associated to each stimulus were equal across interval type (i.e., fifths or tritones) and pitch type (i.e., high or low). Moreover, the ratio between Standard (80%), and Deviant (20%) trials was kept equal across runs.

#### A5 Bayesian Perceptual Surprise Computation

We computed the Bayesian surprise for each trial under a Beta-Binomial model of Bayesian learning of the stimulus probabilities ^12,25,54,55^. The model assumes that the brain implements a sequential learning procedure starting from an uninformative prior, updates this prior according to the subsequent observations, and computes Bayesian surprise as the Kullback-Leibler divergence between prior and posterior. Following Ostwald and colleagues ^25^, we use a variant of this model that assumes an exponential forgetting of stimuli which are observed in the distant past ^12,64^. Formally, we assume that the probability of observing a low (x = 0) or high (x = 1) intensity stimulus at a given trial is described by a Bernoulli distribution with parameter µ ∈ [0, 1]:

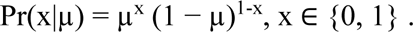

The true value of µ is unknown by the subject, and the initial uncertainty is modelled by means of an informative prior of type Beta, whose density function is uniform on the unit interval:

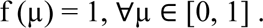

On each trial, the prior is sequentially updated according to the observed data likelihood to form a posterior distribution over µ. After N trials, let x^N^ denote the sequence of observed stimuli

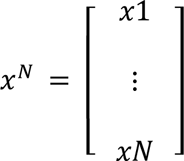

where x_i_ ∈ {0, 1} for each *i* = 1, …, N. Under standard Bayesian learning, the posterior over µ is computed as follows. The probability of observing a x^N^ when the true parameter is µ is:

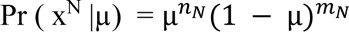

Bayes rule implies that the posterior over µ is a Beta distribution with density

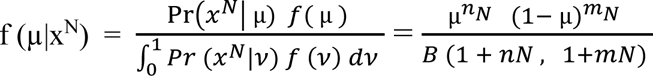

where n_N_ = |{i : x_i_ = 1}| is the number of high-intensity stimuli, m_N_ = N – n_N_ is the number of low-intensity stimuli and B is the Beta function. In order to account for a forgetting dynamic, instead of using the accumulative stimulus counts n_N_ and m_N_ in previous formula, the model employed here weights past observations according to an exponential function. Define the weighted stimulus counts 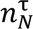 and 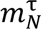 by

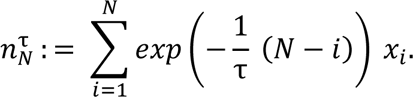

And

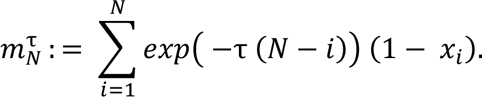

τ ≥ 0 is a parameter governing the forgetting dynamics: for τ = 0 we have 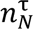, 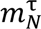 = n_N_, m_N_ whereas increasing the value of τ implies that past observations are weighted less and less. This results in a posterior density given by

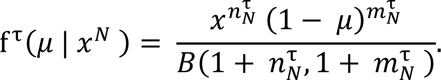

Finally, the model quantifies the degree of learning as the Bayesian surprise, or Kullback– Leibler divergence, between the prior and posterior distribution over µ after a given trial. Let x denote the full sample of observed stimuli. The Bayesian surprise after the N-th trial is given by:

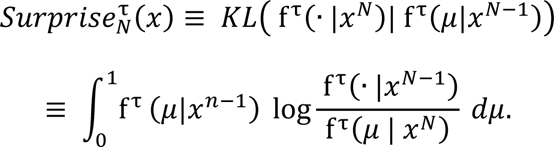

Due to the use of conjugate priors, the Kullback–Leibler divergence can be evaluated analytically, which significantly simplifies the computation ^64^.

### B Data Analyses

#### B1 Analysis in the time domain (ERP) and Mismatch Responses (MMRs)

To compare newborns’ neural response in sensory processing of fifth consonant and tritone dissonant intervals we computed the mismatch response analysis (i.e., MMR). More specifically, we computed the MMR by subtracting the average ERP response elicited by Standard intervals to the average ERP response elicited by Deviant Intervals for each newborn and each condition (i.e. Fifth and Tritone chords ^13^). Crucially we performed the analysis considering only the last interval of the standard sequence (i.e. the one before deviant trials) so as to match the number of standard and deviant trials (N = 52 per run ^12,25^).

##### Group level statistical analyses

To compare the MMR elicited by Fifth and Tritone intervals we performed a whole brain fully data-driven analysis without any aprioristic assumption by computing a point-by-point t test on differential MMR (Fifth vs Tritone). We performed a point-by-point t test ^65^, with cluster size-based permutation correction for multiple comparisons based on temporal consecutivity and spatial adjacency (1,000 permutations; alpha level = .05; percentile of mean cluster sum = 95; minimum number of adjacent channels = 3) on differential MMR (Deviant–Standard). The test compared single subjects’ MMR amplitudes for Fifth and Tritone chords at each time point, for each channel separately. This allowed us to identify time clusters containing mismatch detection responses (Deviant– Standard) that significantly differed between Fifth Consonant and Tritone Dissonant intervals.

Additionally, we performed a “single-point” traditional analysis widely used in newborns studies ^52^, comparing the mean amplitude (Fifth vs Tritone) recorded in a time window and scalp distribution of interest for the research proposal. More specifically, single newborns’ MMR registered on single channel were entered in a group-level analyses to test for possible differences in MMR elicited by fifth consonant and tritone dissonant chords. Based on previous literature ^17,22,40,52^ on newborns MMR, we calculated the mean amplitude within the 0.2-0.45 s range over frontocentral electrodes (FPz, Fz, FP1, FP2, Cz, CP1) for each condition separately (Fifth and Tritone). We then performed a two-tailed t-test between the mean amplitude extracted on the frontocentral electrodes corresponding to the Fifth and Tritone conditions.

##### Group level ANOVA

Finally, we computed a 2 x 2 group-level ANOVA on the event-related waveforms of each participant. Specifically, we set consonance (consonant and dissonant level) and deviancy (deviant and standard level) as a two-level within factor. The statistical analysis corrects for multiple comparisons using a permutation-based correction (1,000 permutations; alpha level = .05; percentile of mean cluster sum = 95; minimum number of adjacent channels = 3).

#### B2 Correlation with Bayesian surprise values

The preprocessing steps in the present analysis were identical to the MMR analysis, with the only substantial difference that all trials must be maintained to compute the correlation between amplitudes and the Bayesian Surprise index. Therefore, we excluded participants with less than 80% trials that survive to the artifact correction. Four participants were excluded from the correlation analysis after visual inspection. However, the remaining sample (N = 18) displays an ERP response to the roving paradigm identical to that of the original sample (N=22), as demonstrated by a significant difference in the polarity of the MMRs elicited by consonant and dissonant intervals (see the Results’ section below for further details). First, a point-by-point trial-by-trial correlation analysis between preprocessed epochs (i.e. 864 for each condition) and the matrix of Bayesian surprise values corresponding to single trials (i.e. 864 for each condition) ^12,31,65,66^ is performed. The analysis computed, for each participant, for each EEG channels and for each condition separately (i.e. Fifth consonant and Tritone dissonant intervals), the correlation between trial-by-trial fluctuations of the EEG traces and the bayesian surprise value. The outcome of the correlation analysis was two 1 s-long (from .2 s preonset to .8 s postonset) time series of r-values for each channel and for each subject.

##### Group level statistical analyses

To verify that EEG amplitudes reflected the bayesian updating as measured by the Bayesian Surprise index, we performed a point by point t-test on each r-values time series (Fifth and Tritone separately) against the constant zero. The two 1 s-long (from .2 s preonset to .8 s postonset) time series of r-values for each channel, for each subject and each condition (Fifth and Tritone) separately, constituted the input for the subsequent group-level two-tailed point-by-point t test with permutation-based correction for multiple comparisons (1,000 permutations; alpha level = .05; percentile of mean cluster sum = 95; minimum number of adjacent channels = 3) against the constant zero. The test compared single subjects’ (N = 18) correlation coefficients, for each condition separately (Fifth and Tritone) against the constant zero at each time point.

Additionally, we compared the mean amplitude of the r-values time series in the 0.2 – 0.45 s range (MMR time interval of interest), over the frontocentral electrodes, for each conditions separately, against the constant zero. Previous studies suggest that mismatch negativity and Bayesian surprise correlation’ peaks share a common time window and scalp distribution, therefore we extracted the mean r values in the same cluster of electrodes and in the same time window of the MMR (FPz, Fz, FP1, FP2, Cz, CP1). We then performed a One Sample t-test to compare mean r values against the constant zero for each condition (Fifth and Tritone) separately.

Furthermore, As in the case of the MMRs analysis, to compare the differences between the r values time series elicited by consonant and dissonant intervals (Fifth vs Tritone), we perform both a point-by-point t test on differential r values time series (Fifth vs Tritone) and a t-test on single subjects’ mean r value.The time series of r-values for each channel and for each subject constituted the input for the subsequent group-level two-tailed point-by-point t test with permutation-based correction for multiple comparisons (1,000 permutations; alpha level = .05; percentile of mean cluster sum = 95; minimum number of adjacent channels = 3). The test compared single subjects’ (N = 18) correlation coefficients, for each condition (Fifth vs. Tritone) at each time point. This allowed us to identify time clusters containing r values that significantly differed between Fifth consonant and Tritone dissonant intervals. We finally compare the mean of the r values time series previously calculated (0.25 – 0.4 s range and over frontocentral electrodes) by computing a two tailed t test (Fifth vs. Tritone). The test compared single subjects’ (N=18) mean r values time series, for each condition (Fifth vs. Tritone).

### C Results

#### C1 ERP Results (MMRs)

The whole brain fully data-driven point-by-point t test performed on mismatch detection responses registered on Fpz (Fifth vs. Tritone) revealed two significant time clusters: the first one centred on the average MMR waveform peak (0.165 – 0.245 s; Figure 2 panel C); the second one occurring later at 0.26 –0.546 s post onset. As expected, MMR waveforms elicited by Fifth and Tritone intervals were significantly different (T = 2.89; p = 0.008 on Fpz). Results were comparable among fronto-central electrodes; the significant cluster corresponding to the MMN extended over Fp1, Fpz, Fp2, F7, F3, Fz.

The two tails paired sample t-test performed on mean amplitudes over the frontocentral electrodes revealed a significant difference between the consonant and dissonant mismatch response, further confirming the point-by-point results (T = −2.22; M = −1.49; SD = 0.67; P = 0.028).

The 18 newborns tested for correlation analysis showed a significant difference in MMRs elicited by consonant and dissonant intervals, identical to the original sample (N = 22). More specifically, the point-by-point t test comparing MMRs elicited by consonant and dissonant intervals revealed a significant cluster in the time interval between 0.26 and 0.48 s among the frontocentral electrodes: Fp1, F7, F3, Fz, Pz (T = 4.61; p = < 0.001; on Fz where the difference is maximal).

##### Group level 2 x 2 ANOVA

The point-by-point ANOVA revealed a significant effect only of the interaction between consonance and deviancy, reflecting the results on mismatch responses described above. More specifically, the statistical comparison revealed a cluster of significance (i.e., effect is maximal on Fpz: F = 3.51; p = 0.009) of interaction (consonance x deviancy) in the time interval between 0.16 and 0.55 s in frontocentral electrodes (Fp1, Fpz, Fp2, F7, F3, Fz).

#### C2 Trial-by-Trial Correlation with Bayesian Surprise

The correlation analysis between single trial amplitudes and Bayesian Surprise indicated that r values peaked at 0.1 s (corresponding to the MMN peak latency; Figure 2 B in the main text) for both Fifth and Tritone intervals. Subsequent fluctuations in r values peaked at 0.3 s and 0.4 s postonset corresponding to the P3 and the N400 components (see Figure 2 B in the main text). These results confirmed our prediction, indicating that MMR best indexes Bayesian perceptual learning in our study.

The whole brain point by point t-test against the constant zero revealed that both Fifth consonant and Tritone dissonant time series of r-values resisted to the 1000 permutation and were significantly different from zero. More specifically, the time series of r-values elicited by fifth consonant interval were significant in the time window between 0.1 and 0.4 s postonset over frontocentral channel (F4 and P3; p = 0.004 and T = −3.25 on F4, where the difference in maximal) while the r values time series elicited by tritone intervals were significantly different in the time window around 0.6 s over frontocentral channel (Fpz, F7, F3, Fc1 and Cz; p = 0.008 and T = 2.98 on Fpz where the difference is maximal). The One sample t-test performed on the mean amplitude extracted over the frontocentral electrodes (FPz, Fz, FP1, FP2, Cz, CP1) comparing Fifth consonant (T = −2.00; M = −0.006; SD = 0.03; P = 0.04) and Tritone dissonant (T = 3.42; M = 0.01; SD = 0.008; P = 0.003) conditions against the constant zero for each subject reveal a significant difference between the consonant and dissonant r-values time series, further confirming the point by point results.

The point-by-point t test performed between the r values time series of fifth and tritone conditions revealed that the two r-values waveforms (Fifth vs. Tritone) were significantly different in the time interval between 0.1 and 0.4 s (Corresponding to the latency of MMR) over frontocentral electrodes (Fpz, Fp2, F4, Fc1, Fc2). More specifically, the first significant cluster is found in the time interval between 0.06 and 0.13 s over Fp2 (p = 0.006; T = 3.02) while the second larger cluster is found in the time interval between 0.28 and 0.4 s over Fpz, Fp2, F4, Fc1, Fc2 (p = 0.006 and T = 3.1 on Fpz where the difference is maximal). The two tails paired sample t-test performed on the mean amplitude extracted on the frontocentral electrodes comparing Fifth consonant and Tritone dissonant conditions (Fifth vs Tritone) reveal a significant difference between the consonant and dissonant r values time series (T = −4.09; M = −0.017; SD =0.004; P < 0.001), further confirming the point-by-point results.

